# A *de novo* DNA Sequencing and Variant Calling Algorithm for Nanopores

**DOI:** 10.1101/019448

**Authors:** Tamas Szalay, Jene A. Golovchenko

## Abstract

The single-molecule accuracy of nanopore sequencing has been an area of rapid academic and commercial advancement, but remains insufficient for the *de novo* analysis of genomes. We introduce here a novel algorithm for the error correction of nanopore data, utilizing statistical models of the physical system in order to obtain high accuracy *de novo* sequences at a range of coverage depths. We demonstrate the technique by sequencing M13 bacteriophage DNA to 99% accuracy at moderate coverage as well as its use in an assembly pipeline by sequencing λ DNA at a range of coverages. We also show the algorithm’s ability to accurately classify sequence variants at far lower coverage than existing methods.

---

DNA sequencing has proven to be an indispensable technique in biology and medicine, largely due to developments in low cost and high throughput 2^nd^ and 3^rd^-generation technologies^1,2^. Despite these advances, however, most existing sequencing-by-synthesis techniques remain limited to short reads using expensive devices with complex sample preparation procedures^3^.

Initially proposed two decades ago by Branton, Deamer, and Church^4^, nanopore sequencing has recently emerged as a serious contender in the crowded field of DNA sequencing. The method uses a small trans-membrane pore whose narrowest constriction is just wide enough to allow single-stranded DNA to pass through (Fig. 1a). An applied voltage across the membrane sets up an ionic current and electrophoretically draws the DNA into the pore. This current is monitored to measure the changes in conductance caused by the presence of DNA. An enzymatic motor such as a polymerase^5^ or helicase is used to ratchet the strand through the pore one base at a time and the resulting changes in ionic current can be used to deduce the sequence.

**Figure 1.**
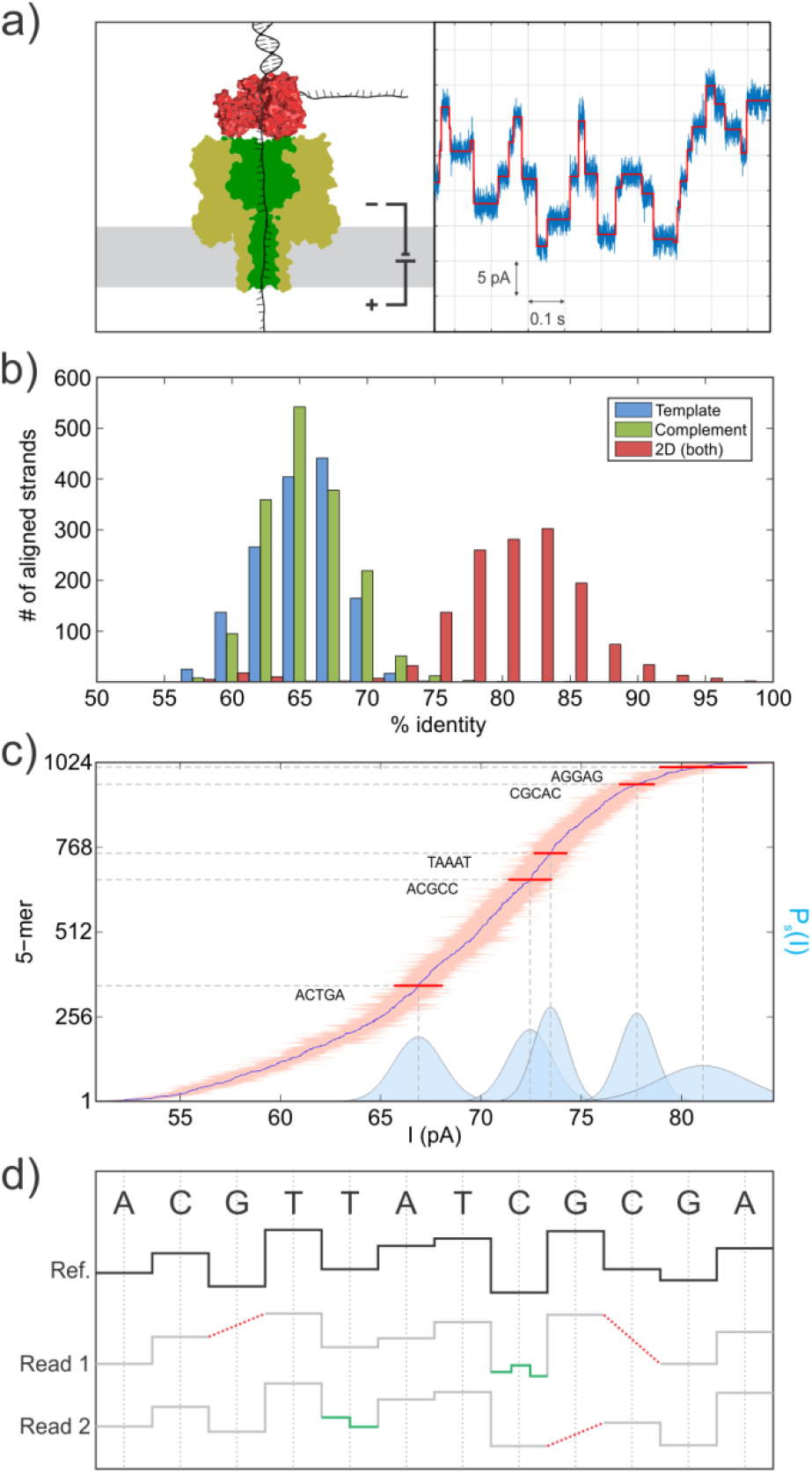
a) Illustration of the DNA-enzyme complex captured in a nanopore (left). The base-by-base processive behavior of the ATP-fueled ratcheting enzyme leads to the depicted ionic currents (right) which are discretized to facilitate subsequent analysis (red line). b) Summary analysis of a sequencing run of M13mp18 DNA on an Oxford MinION device demonstrating the available depth of coverage at moderate accuracy. Each data point represents an entire M13 DNA molecule. c) A plot of the mean currents and standard deviations of the 1024 distinct 5-mer sequences, with the full Gaussian distributions of a few example 5-mers shown in blue. d) Depiction of the alignment issues caused by possible detection errors in multiple reads (grey) against the expected ideal current levels (black), including missed levels (red) and extra levels (green).

Nanopore research groups have recently demonstrated the feasibility of obtaining long-read data with quantifiable accuracy^6^, and Oxford Nanopore Technologies has released their 2,048-nanopore USB-powered MinION sequencer to a public open access program^7–9^. The device makes use of integrated current amplifiers and consumable flow cells along with biochemical sequence preparation kits in order to collect tens to hundreds of megabases of data in a single run. These developments have enabled nanopore sequencing to produce data at high coverage and moderate accuracy (Fig. 1b), while also encouraging the creation of freely available tools and techniques for subsequent analysis^10–12^. Such long-read data have been used as a scaffold to aid in the assembly of shorter, more accurate reads^8,13^; however, few techniques exist for combining low accuracy reads directly^14^. Here we show that the latent information in the ionic current data from multiple reads can greatly increase the accuracy when coupled with proper statistical modeling of the underlying physical system.

The dominant source of uncertainty in nanopore sequencing is the simultaneous influence of multiple adjacent nucleotides on the ionic current signal. It has previously been shown that up to 5 bases influence the instantaneous current^15,16^, increasing the number of distinct current levels from the ideal of 4 up to as many as 1024 and thus having a deleterious effect on the signal-to-noise ratio for base determination (Fig. 1c). The difficulty of extracting the sequence is further compounded by the stochastic behavior of the DNA, enzyme, and nanopore complex, which can lead both to missing and extra current levels, as illustrated in Fig. 1d. The skipped levels can be caused by the enzyme randomly ratcheting past a particular base too quickly to detect, and, as a result, the discretized form of the data (Fig. 1a, red line) will have that particular level omitted. Fluctuations or conformational changes can also lead to sudden jumps in conductance that could easily be mistaken for actual level transitions even though the enzyme stays on the same base, and certain enzymes can even exhibit random backwards motion^17^. These confounding factors lead to a problem of alignment: there is no longer a one-to-one correspondence between the detected sequence of current levels and the true sequence of bases. The large number of possible mappings between levels and bases thus results in many more actual sequences that could potentially produce a particular set of observed data.

We introduce a simple statistical model to capture the above effects, composed of both the probabilities of observing a current level given a 5-mer sequence (Fig. 1c) and the probabilities of skipping or staying on a given current level (Fig. 1d, Supplementary Fig. S1). Then, given a sequence, a set of observed current data, and an alignment that maps the discretized levels to the positions in the sequence, these probabilities can be multiplied together to compute a total observation likelihood for that sequence. The best sequence is defined as the one that results in the highest likelihood over the observations being considered, whether from one or multiple reads.

Although techniques exist for finding the maximum likelihood sequence using the dynamic programming^18^-based Viterbi algorithm^19,20^, they are limited to analyzing only one or two reads at a time due to the computational complexity of simultaneously aligning the levels and finding the sequence (Supplementary Note 1). Other methods such as dynamic time warping^6,21^ can align many sets of levels to one another without prior knowledge about the sequence, but in doing so, they sacrifice valuable information contained in the statistical model. In order to reliably surpass the accuracy obtained from single-molecule reads (Fig. 1b), a new approach is thus needed.

The POISSON (Plural Observation-aligning Iterative Sequence Space Optimization for Nanopores) algorithm presented here is designed to accurately determine the *de novo* sequence using the discretized ionic current data from an arbitrary number of independent nanopore reads of the same region of DNA, including partial or reverse complement reads. It does this by iteratively finding the sequence that maximizes the total observation likelihood for all of the reads according to the statistical model. Optimizing the sequence in this iterative way requires searching through the vast space of all possible sequences of a given length; POISSON limits the search to sequences that could plausibly fit the data by repeatedly testing and making only those local changes that are expected to improve the observation likelihood.

The components of the algorithm are illustrated in Fig. 2a. The process starts with a candidate sequence considered to be an initial best guess for the optimization routine and a number of observed nanopore reads that cover some of the region over which we are optimizing. A natural choice for the initial guess is one of the read’s single-molecule sequences computed via existing Viterbi methods (e.g. as provided by Oxford Nanopore). This guess is then gradually improved by introducing artificial mutations into the candidate sequence; these mutations are drawn from alternate versions of the candidate sequence, generated by a modified single-molecule Viterbi algorithm that deliberately introduces some randomness (Supplementary Note 2). Although these alternate sequences globally fit the data slightly worse than the best candidate, they may contain a short region (anywhere from one to dozens of bases) that has a higher sequence-aligned likelihood in that region compared to the current best candidate (Fig. 2a, iii, shaded). This stretch of sequence is then locally substituted into the best candidate’s sequence and the full observation likelihood recalculated, and the mutated sequence is kept if this likelihood exceeds the current best. Once all such mutations have been exhaustively tested, new alternates are generated and the procedure repeats until no more changes are found. This technique makes sequence space optimization feasible by screening for likely mutations and thus avoiding a prohibitively dense search over all possible sequences.

**Figure 2.**
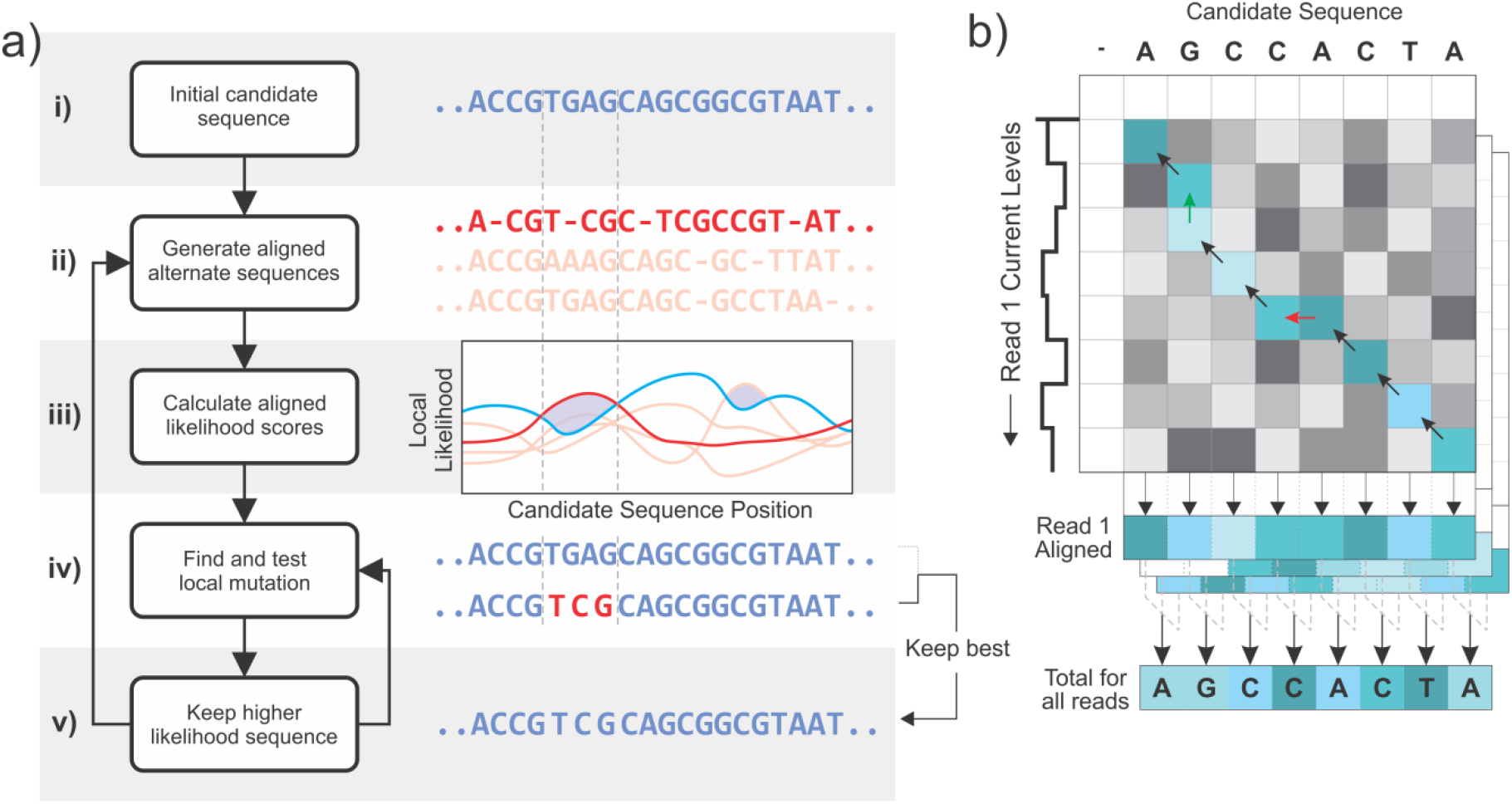
a) A high-level overview of the POISSON algorithm. The process starts with multiple nanopore reads and a single candidate sequence shown in blue (i). Aligned alternate sequences from which mutations are drawn are shown in red with a specific one highlighted (ii). The alternates must be similar enough to the candidate that their sequences can be aligned, but otherwise vary throughout their full length. The plot contains the aligned local likelihood scores across all reads (calculated as in 2b) for the candidate and each alternate (iii). Local regions where an alternate’s likelihood score exceeds the candidate’s are shaded and correspond to a short region where the alternate sequence is likely more accurate than the best candidate. The alternates’ local sequence from this region is substituted into the best candidate and the full maximum likelihood score is recomputed (iv), at which point the higher-scoring sequence becomes the new best (v). The vertical dotted lines denote a specific mutation taken from the highlighted red strand in parts (i) through (iv). b) The maximum likelihood over all possible alignments of a single read and sequence is computed using a Smith-Waterman^22^-style dynamic programming approach. The values of the matrix elements are the maximum likelihood over all valid paths to that element while the lightness of the color represents the local probability of that particular sequence/current pair according to the statistical model. The maximum likelihood score is the highest value in the whole matrix and the full best alignment can be found by backtracking through the matrix as illustrated by the arrows. The local probabilities of each aligned read can be mapped back to the sequence and combined (bottom), at which point they can be compared to the scores from other sequences (2a, iii, comparison between red and blue lines).

The details of the likelihood calculation are illustrated in Fig. 2b. The matrix depicts the alignment of a single read with a candidate sequence and yields the maximum likelihood (over all possible alignments) of observing that read given the sequence. The calculation of each cell in the matrix proceeds in a similar manner to canonical alignment algorithms (dynamic time warping^21^ or Smith-Waterman^22^): the row and column coordinates of a single cell represent the indices of the current levels and the bases, respectively, and the value contained in that cell is the maximum likelihood over all possible paths that can be taken to reach that cell. The cells are filled in using the likelihood values from the surrounding three cells directly above and to the left, as they correspond to the different possible cases of enzymatic motion according to the statistical model (e.g. a step from the left is a skip, shown in red, as the sequence advances without a corresponding current level). The resulting likelihood value of the new matrix cell is the maximum likelihood out of all of these starting cells multiplied by the probability of the new sequence/observation pair and the skip/stay probabilities, if applicable (Supplementary Note 3). At the end of the procedure, the highest likelihood in the entire matrix is then the maximum likelihood over all alignments, and the best alignment itself can be obtained by tracing the matrix backwards (shaded blue, arrows) from the maximum. A separate matrix is calculated for each read and the resulting likelihoods are then multiplied together to get the total observation likelihood; the computation time hence scales linearly with the coverage depth.

The algorithm’s ability to obtain *de novo* sequence information using MinION nanopore data is shown in Fig. 3a. The data are taken from M13mp18 digested by EcoRI, yielding identical double-stranded fragments 7,249 bases long, and from *λ* phage DNA sheared to approximately 8 kb fragments (Supplementary Fig. 5). After preparing the molecules as recommended (see Methods), we ran a 24-hour sequencing protocol. The initial sequence analyses were done using Oxford Nanopore Technologies’ cloud-based Metrichor service, which computed the sequence corresponding to each detected molecule separately. The models (internally trained on the *E. coli* genome^23^) used by Oxford to map 5-base sequences to observed current levels were also stored, and these models were used in this work without modification along with the offsets and scaling provided. The additional skip and stay parameters for the model were trained using the results of a sequencing run of identical 3.6kb fragments of DNA that ship with the MinION device for calibration (see Methods).

**Figure 3.**
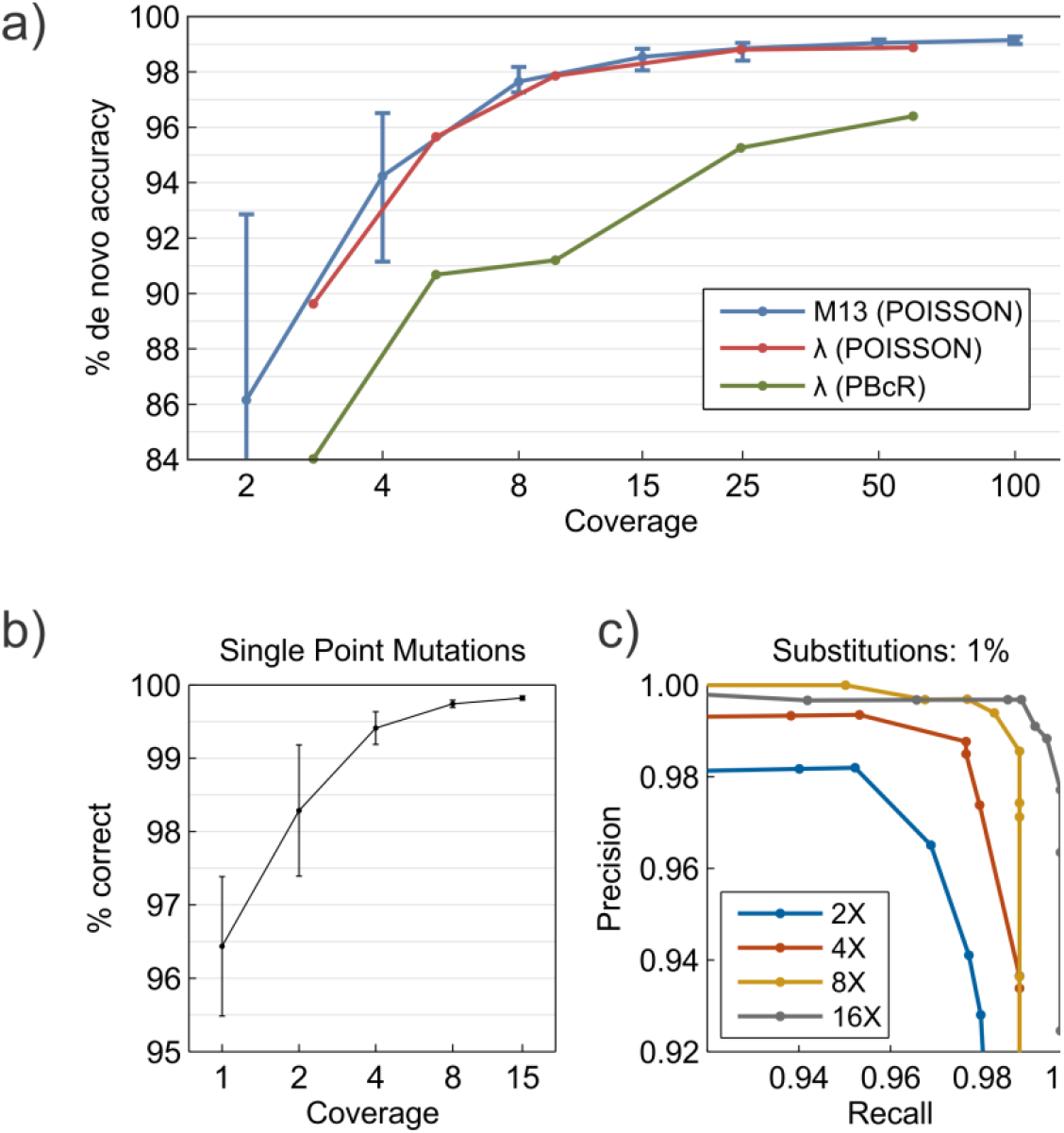
a) Accuracy results of running POISSON on nanopore data from M13 and λDNA to obtain *de novo* sequences. For M13, error bars indicate the upper and lower bounds for accuracy across 20 random subsets at the given coverage. The green line is the result of error correction and assembly with PBcR using only the 2D basecalled sequences; the red line shows the improvement when POISSON is used to error-correct with the raw data. b) Fraction of single-base variants of M13mp18 correctly called as a function of coverage. Variant sequences were generated by computationally making every possible insertion, deletion, or mutation in the original sequence of M13. A correct call is defined as the original M13 sequences’ likelihood being larger than the variant in question. Error bars denote the deviation across 20 random subsets of molecules. c) Variant calling performance of POISSON on substitution mutations introduced in M13 at a higher frequency of 1%, at a range of coverages. Precision and recall denote the probabilities of false positives and negatives, respectively. The maximum *F-*score accuracy shown is 99.1% at 16X coverage.

The *de novo* sequence accuracies of M13mp18 are shown in Fig. 3a as a function of single-molecule coverage, with each molecule consisting of both a template and complement read. The error bars represent the minimum and maximum accuracy obtained from trials using different random subsets of molecules at the specified coverage. We emphasize that no information about the true sequence was used, nor any information (e.g. statistics or fit parameters) from molecules outside of those included in a single trial. Due to the higher levels of coverage needed to reach a practical 99% accuracy, small DNA samples might require the use of PCR, in which case the mutation scoring would be modified to discard the lowest-scoring strands so as to account for the errors introduced during replication.

We also demonstrate the use of POISSON in a standard genome assembly pipeline by assembling *λ* DNA from nanopore reads at a range of coverages (Fig. 3a). Our technique is based on the Hierarchical Genome Assembly Pipeline (HGAP)^14^ developed for Pacific Biosciences sequence data, consisting of three stages: fragment error correction, assembly, and assembled genome refinement. When the existing pipeline is run on MinION sequence data directly, the accuracy is limited to around 96% (Fig. 3a, green line); however, replacing both the existing fragment error correction and refinement stages with POISSON (see Methods) increases the accuracy to 99% (Fig. 3a, red line). Importantly, long sequences are split into smaller sections for parallel processing, making the technique inherently scalable to larger genomes (see Supplementary Note 7 for a discussion of execution time).

Another benefit of POISSON is that it easily enables sequence variant comparison, as shown in Figs. 3b and 3c. It is often necessary to distinguish between known single nucleotide variants (SNV) occurring at a low density (approx. 1 SNV per 2 kb in humans^24^). As a test, we took the actual M13mp18 sequence and computationally mutated it in one position (replacing, inserting, or deleting a single base) to generate an SNV of the original sequence. The observation likelihood feature of POISSON was then used to compute the likelihood score of both the original and mutated sequences in order to call the correct variant as the higher of the two likelihoods. As can be seen in Fig. 3b, even at low coverages POISSON accurately identifies the correct unmodified sequence as having a higher likelihood score. This variant calling feature outperforms *de novo* sequencing at lower coverages because the vast majority of single nucleotide-mutated sequences would yield a considerably different current signature, making them easy to distinguish.

We also compare our approach to previous sequence-based variant calling analyses using nanopore data^12^ (Fig. 3c). Substitution errors are introduced into the M13mp18 sequence at a rate of 1%, and we then attempt to correctly identify them using POISSON and recover the original sequence. By setting a threshold on the difference in observation likelihood when calling each base variant, we study the probability of false positives and false negatives (referred to as precision and recall, respectively) at a range of coverages, in order to find the maximum classification accuracy. We find a peak *F*-score accuracy of 99.1% at a coverage of only 16X, demonstrating similar performance to previous results^12^ but at an order of magnitude lower coverage.

The results in this work show that multiple nanopore reads can be combined to reach accuracies over 99%, compared to the approximately 85% seen with single molecules^7^, and many further improvements are expected both in the nanopore biochemistry and the physical models capturing their behavior. In particular, the inclusion of current level durations will be necessary in dealing with homopolymer DNA regions, which we have found are responsible for over half of the errors at 99% accuracy (Supplementary Note 5). The model can also be extended to detect base methylation, the identification of which requires an estimated 5-19 repeated reads^25,26^. Other improvements are possible through better control of enzymatic ratcheting^27^ or the inclusion of a wider variety of pore mutants^28–30^ to obtain pore-specific current data on the same sequence. The POISSON algorithm was designed to be flexible in handling such modifications with far fewer constraints than Viterbi-based approaches, and we have made all code publicly available at http://github.com/tszalay/poisson in the hope that others will make use of it as a framework for future development.

## ACKNOWLEDGMENTS

We would like to thank E. Brandin for molecule preparation, D. Branton for obtaining MinION sequencers, S. Fleming for helpful algorithmic discussions and Fig. 1a, and A. Kuan and M. Burns for feedback on this manuscript. The computations in this paper were run on the Odyssey cluster supported by the FAS Division of Science, Research Computing Group at Harvard University, and the work was supported by the National Institutes of Health Award No. R01HG003703 to J.A. Golovchenko and D. Branton.

## AUTHOR CONTRIBUTIONS

T.S.: algorithm development, data analysis and interpretation, writing of manuscript, J.A.G.: data analysis and interpretation, writing of manuscript.

## ONLINE METHODS

### M13 Restriction Digest

Four micrograms of M13mp18 RFI (New England Biolabs, cat. no. N4018S) DNA were digested with EcoRI restriction enzyme in a 100 microliter reaction volume for 2 hrs at 37C, and then heated for 30 min at 65C to inactivate the enzyme. The digested DNA mix was applied to Micro Bio-Spin 30 chromatography columns (Bio-Rad Laboratories Inc., cat. no. 732-6223) equilibrated with molecular grade water, centrifuged according to the manufacturer’s instructions, and a column flow-through volume containing one microgram of desalted DNA was adjusted to 85 microliters with water for subsequent library preparation.

### Lambda DNA Shearing

Lambda genomic DNA for sequencing was prepared by shearing the DNA using the Covaris g-TUBE (Covaris, Inc., cat. no. 520079) to generate targeted DNA fragment sizes of approximately 8kbp. One microgram of Lambda DNA (New England Biolabs, cat. no. N3013S) in a total volume of 80 microliters water was applied to a Covaris g-TUBE and centrifuged, according to manufacturer’s instructions, for one minute at 7,200 RPM in a Heraeus Biofuge fresco (Kendro Laboratory Products) with temperature set at 30C. Five microliters of DNA CS (ONT) was added to the sheared DNA and the sample prepared for sequencing following the Oxford Nanopore MAP Genomic DNA Sequencing Kit protocol (ONT prod. Code SQK-MAP003).

### Sequence Library Preparation

End repair of the digested DNA samples was performed using NEBnext End Repair module (New England Biolabs, cat. no. E6050S) by adding 10 microliters of reaction buffer and 5 microliters of enzyme mix and incubating at room temperature for no longer than 30 minutes. To purify the end-repaired DNA, 100 microliters (1X volume) of Agencourt AMPure XP beads (Beckman Coulter Inc., cat. no. A63880) were gently mixed in, incubated at room temperature for 15 min, and the beads separated from the reaction supernatant using a magnetic rack. DNA was eluted from the beads in 25 microliters of 10 mM Tris-HCl pH 8.5 and dA-tailing performed at room temperature for 60 min by adding 3 microliters of reaction buffer and 2 microliters of enzyme supplied in the NEBNext dA-Tailing module (New England Biolabs, cat. no. E6053S).

The Oxford Nanopore MAP Genomic DNA Sequencing Kit (ONT prod. Code SQK-MAP004 for M13, SQK-MAP003 for lambda) was used to further prepare the DNA samples for sequencing. Kit reagents were thawed and stored on ice (HP adaptor, adaptor mix, and fuel mix), or at room temperature (2X wash buffer, elution buffer, and EP buffer) before use. Following the manufacturer’s protocol, a ligation step was performed by combining (in order) 30 microliters of dA-tailed DNA, 8 microliters of water, 10 microliters of Adaptor mix, 2 microliters of HP adaptor, 50 microliters of Blunt/TA Ligase Master Mix (New England Biolabs, cat. no. M0367S), and incubating the mixture at room temperature for 10 minutes. Ten microliters of His-Tag Dynabeads (Life Technologies, cat. no. 10103D) were washed twice with 250 microliters of 1X wash buffer and suspended with 100 microliters of 2X wash buffer. This volume of washed His-Tag beads was added directly to the ligation mixture, incubated for 5 min at room temperature, and the beads collected using a magnetic rack. Once separated from the reaction supernatant, the pelleted beads were carefully rinsed twice (without suspension) using 250 microliters of 1X wash buffer. All wash buffer was subsequently removed and the DNA Library eluted from the beads with 25 microliters of elution buffer. Prior to sequencing on the MinION, 140 microliters of EP buffer was added to a 6 microliter aliquot of the DNA library, followed by 4 microliters of fuel mix and thorough mixing.

### Data Acquisition

Data was acquired with an Oxford MinION device connected to a sufficiently capable computer (Windows 7; MinKNOW software; USB3; SSD; i7 processor). A new version R7.3 flowcell was used for each run; the flowcell was initially primed with two injections of 150 microliters of EP Buffer, spaced 10 minutes apart, after which the MinKNOW software was used to measure pore quality and activity. The 150 microliters of previously prepared sequencing mix was then loaded into the MinION flowcell as specified by the manufacturer. A 24-hour sequencing protocol was selected in the software and the device was allowed to run until all pores and sample were exhausted.

### Data Analysis

Initial data analysis was performed by Oxford Nanopore Technologies’ cloud-based Metrichor service, which runs proprietary versions of the Viterbi algorithm similar to those described in the Supplementary Information. The service uploads the basic HDF5 (‘.fast5’) files generated by the sequencing software and returns the files with additional processed data appended. A number of these analyzed features were used in the present work: the parsing of each molecule to separate the current levels corresponding to the template and complement strands in each double-stranded molecule (and the removal of hairpin/adapter levels); the Oxford-trained statistical models used to map 5-mers to current levels; the offset and scaling between the molecule and model current levels; estimates of the skip and stay probabilities per molecule; and the two 1D Viterbi and one 2D Viterbi-computed sequence for each molecule. The details of the extraction and use of each of these features is well-documented in Supplementary Note 6 and the provided source code. Only the models and scaling are required; the Viterbi sequences were merely used to help speed up convergence of the algorithm.

### Model Training

The skip, stay, and insertion parameters of the model were trained using a sample of ONT-provided “DNA CS” (calibration sample) prepared in the same manner as the M13mp18 post-digest using the material supplied with the SQK-MAP004 kit. Of the 907 molecules sequenced, a subset of 20 was selected at random and *de novo* sequencing performed using POISSON. The parameters were arbitrarily seeded to 5% for skips/stays and 2% for insertions (Supplementary Note 3), then randomly perturbed and the change kept if they increased the resulting DNA CS sequence accuracy. After a few hundred iterations, the parameters had converged and no improvements were found; these were the parameters used in the final analysis used to generate Fig. 3. The authors acknowledge that more precise means of training the parameters are possible, but it was found that the *de novo* accuracies are fairly insensitive to small (<20%) changes in the parameters, and as a result the present method was found to be sufficient.

### Accuracy Calculation

When the term “accuracy” was used to refer to a DNA sequence relative to the M13mp18 reference (Figs. 1b, 3a), this accuracy was calculated as follows: first, the optimal alignment between the candidate sequence and a reference was found using Matlab’s swalign function (Smith-Waterman alignment) using default parameters, which uses a match score of +5, a mismatch penalty of -4, and a gap/extend penalty of -8. The alignment is assumed to be good enough to cover all of the candidate sequence unless otherwise noted. Next, the accuracy was computed as the number of matching bases divided by the total number of bases in the alignment, defined as matches + mismatches + insertions + deletions. For the variant calling and *de novo* sequencing trials of M13, the accuracy was calculated in the region covered by at least 75% of the strands, to compensate for end-trimming in the data. In the case of DNA, the accuracy reported is % identity as calculated by MUMmer^31^, which was found to not differ significantly from other definitions at the error rates shown.

### M13 *de novo* Sequencing

To reconstruct the original sequence from nanopore reads of M13 bacteriophage DNA, POISSON was executed on a specific number of fragments as shown in the coverage plot of Fig. 3. Suitable fragments were defined as those with double-stranded information available whose 2D basecalls had between 5000 and 8000 bases (compared vs. the full M13 with 7249), in order to filter out partial or repeated molecules. POISSON was then run using each molecule’s own 2D Viterbi sequence generated by Metrichor as the alternate sequence inputs to the algorithm, until no mutations were found, typically taking around 3 iterations (with up to a thousand mutations possible in each iteration). Once this phase completed, alternate sequences were generated as described in Supplementary Note 2, to find likely mutations that did not appear in any of the individual 2D sequences. In each iteration, 12 alternate sequences were generated with identity/similarity to the candidate ranging from 60% to 100%, and these iterations were alternated with the testing of every possible single-base mutation (Supplementary Note 6) to ensure that no mutations were missed; this was repeated at most 5 times and found to sufficiently capture all mutations.

### Lambda Assembly

POISSON was used to error-correct each individual read as follows: first, the 2D-basecalled sequences of all reads were extracted from the returned ONT files. LAST was then used to perform overlap alignment of each read’s sequence to all other sequences, and these alignments were used to seed the POISSON algorithm and improve the accuracy of each fragment sequence. The latest Celera assembler (v8.3) was then used to assemble the corrected sequences into a single contig. All reads were re-aligned with the contig once again using LAST, and corrected in 6000-base fragments which were finally reassembled into a single, high-accuracy sequence. All commands and parameters used can be found in the Supplementary Code. Each point in Fig. 3a is from a single random subset of all reads, and the coverage shown is the theoretical maximum value of (total bases in reads) / 48502. The specifics of the iterations for the correction are the same as that described in the previous section.

